# Health seeking behaviour and cost of fever treatment to households in a malaria endemic setting of northern Ghana: A cross sectional study

**DOI:** 10.1101/486183

**Authors:** Maxwell Ayindenaba Dalaba, Patricia Akweongo, Philip Baba Adongo, Philip Ayizem Dalinjong, Samuel Chatio, Abraham Oduro

## Abstract

**Background:** This study examined the health seeking behaviours and cost of treatment of malaria to households in the Kassena-Nankana district of Ghana.

**Methods:** A cross-sectional household survey was conducted between July and September 2015. Individuals who had an episode of fever or malaria in the past two weeks identified during routine Health and Demographic Surveillance System data collection were selected for the study. Socio-demographic characteristics, treatment seeking behaviours and cost of treatment of malaria were obtained from the patient perspective.

**Results:** Out of the 1,845 households visited, 21.3% (393/1,845) reported to have had an episode of fever or malaria in the past two weeks. Of the 393 people with malaria, 66.9% (263/393) reported taking an antimalarial. About 53.6% (141/263) of the antimalarials were obtained from formal healthy facilities. About 36.1% (95/263) reported to have taken Dihydroartemisinin-piperaquine, 35.4 % (93/263) took Artesunate–Amodiaquine and 21.7% (57/263) took Artemether-Lumefantrine. Only 49.6%% (195/393) of the study participants had their blood sample taken for illness (microscopic or Rapid Diagnostic Test). Only 23.6% (62/263) took antimalarial within 24 hours of the onset of illness. The overall average costs (direct and indirect cost) incurred by households per malaria treatment was GH¢27.82/US$7.32 (range: GH¢0.2/ US$0.05 - GH¢200/ US$52.63). The average cost incurred in the treatment of malaria by those who were enrolled into the National Health Insurance Scheme (NHIS) was GH¢24.75/US$6.51 and those not enrolled was GH¢43.95/US$11.57.

**Conclusions:** Prompt treatment such as treatment within 24 hours of onset of malaria was low. The average costs to households per malaria treatment was GH¢27.82/US$7.32 and the preferred antimalarial was Dihydroartemisinin-piperaquine. There was a positive effect of NHIS enrolment on cost of treatment as the insured incurred less cost (US$5 less) in treatment than the uninsured.

## Introduction

According to the WHO report, there are an estimated 198 million malaria cases and 584 000 malaria deaths worldwide in 2013. The burden of malaria is more severe in sub-Saharan Africa where about 90% of all malaria deaths occur [1]. It is a major cause of poverty and slows economic growth by up to 1·3% per year in endemic countries[2].

Malaria is a significant public health problem in Ghana where it accounts for about 33% of all outpatient attendances and 49% of under five years admissions[3]. Malaria is a preventable, diagnosable and treatable illness. The main methods for the prevention of malaria include Long-Lasting Insecticidal Nets (LLINs), Indoor Residual Spraying (IRS), Intermittent Preventive Treatment for pregnant women (IPTp) and vector control methods such as larviciding [1,4]. The WHO has recommended that all persons from endemic areas with suspected malaria should be examined for evidence of infections with malaria parasites by Rapid Diagnostic Test (RDT) or microscopy. In addition, WHO recommended that uncomplicated malaria should be treated with antimalarial such as Artemisinin-based combination therapies (ACTs), particularly in areas where malaria is endemic and persons should have access to ACTs within 24 hours of onset of malaria [1].The recommended ACT combinations are artemether-lumefantrine (AL), artesunate-amodiaquine (AS+AQ), artesunate-mefloquine (AS+MQ), dihydroartemisinin-piperaquine (DP), and artesunate-sulfadoxine-pyrimethamine (AS+SP). The choice of ACT in a country or region is based on the level of resistance of the partner medicine in the combination [5,6].

Following WHO recommendation, in 2004 Ghana changed her antimalarial drug policy choosing Artesunate-Amodiaquine combination as the first line drug for the treatment of uncomplicated malaria to replace chloroquine. However, the implementation process was confronted with some challenges which relates to adverse drug reactions, safety concerns and absence of other treatment options. It therefore became necessary to review the new policy to address all recognized concerns. A task force reviewed the existing policy and selected additional ACT drugs and dosage forms to accommodate for those who could not bear the Artesunate-Amodiaquine combination. Two additional first line ACTs namely; Artemether-Lumefantrine and Dihydroartemisinin-Piperaquine were selected thus making Ghana to have three official concurrent first line antimalarial treatments [7].

To date, however, the reaction of the health system to multiple alternative first line malaria therapy has not been comprehensively studied. Little is known about access to effective and prompt treatment and preferences for antimalarial drugs for treatment of fever/malaria as well as cost of malaria treatment in the face of multiple first line treatment policy. This study therefore examined the health seeking behaviours as well cost of treatment of malaria to households in the Kassena-Nankana district of Ghana in the face of multiple first line antimalarial drugs.

## Methods

### Study area

The study was carried out in the Kassena-Nankana East and West Districts of the Upper East Region of Ghana. The Kassena-Nankana East is home to the Navrongo Health Research Centre (NHRC) and staff of the Centre conducted the study. The NHRC operates Health and Demographic Surveillance System (HDSS) and has a data base of all individuals and households in the Kassena-Nankana East and West Districts. For the purpose of research, the NHRC refers to the two districts by their former name - the Kassena-Nankana District (KND).

The KND covers an area of about 1,674 square kilometres of land [8]. The area is characterized by a short rainy season and a prolonged dry season from October to March. The mean annual rainfall is about 850mm, with the heaviest usually occurring in August. The total population currently under surveillance is about 152 000 residing in about 32,000 households [8]. Malaria transmission in the KND occurs all year round but there is a distinct seasonal pattern with the peak of transmission coinciding with the period of the major rains (August) and the dry season seeing very low rates of malaria infection[9].

The districts have one district referral hospital located in Navrongo town and 8 health centres strategically located across the district which provide secondary curative and preventive health care. There are 28 Community Based Health Planning and Services (CHPS) compounds/clinics located in various communities and providing primary health care treatment for minor ailments and also carrying out childhood immunizations and antenatal services. There are three private clinics, three pharmacies and over 50 licenced chemical shops in the area.

### Study design

A cross-sectional household survey design was used, and data was collected using quantitative approach. The study was conducted within the INDEPTH-network effectiveness and safety studies (INESS) Phase IV research platform that aimed to assess the effectiveness of new malaria treatments and its determinants in real life health systems[10,11].

### Sample size and data collection

A standardized, rolling, nested, household survey was used, by employing two-week recall period. We assumed conservatively that at least 6% of the population would have had fever event two weeks prior to the interview date and that about 50% of the population with fever gained access to an official point of ACT provision within 24 hours. We accounted for clustering within households by using a design effect of 1.5. We further allowed a 10% drop out rate to achieve a sample size of 1, 845 households.

The data for this study were collected as an additional module within the framework of the Navrongo Health and Demographic Surveillance System round updates. Field workers visited the sampled households between July and September 2015. All members in the selected households with a history of fever in the prior two weeks were interviewed using a structured questionnaire. For children who were less than 18 years of age their adult caretakers were interviewed. Information on socio-demographic characteristics, treatment seeking behaviours and cost of treatment were obtained. Cost of treatment was collected from the patient perspective.

Those who treated with antimalarial drugs were asked the name of the drug and a physical observation of the drugs if it was still in use or if the package was kept. Field workers carried photo images of the variety of antimalarial packages (brand and generic names) available from private and public providers in the district to facilitate identification of treatments received. Types of antimalarial drugs were to be captured on the questionnaire by their generic names. For instance, brands such as Camosunate, Amonate, Coarsucam, to be captured as *Artesunate-Amodiaquine*; Malar-2, Lonart, lumartem, Coartem, Co-Malagon, Lumether as *Artermether-Lumefantrine*; and, P-Alaxin, Eurartesin, Duo-Cotecxin, as *Dihydroartemisinin-piperaquine*.

### Data processing and analysis

Data were double entered into fox pro and transferred to Stata 11.0^©^ for analysis. We used descriptive statistics showing proportions, means and numbers.

Ages of respondents were captured as continuous variables but during analysis were grouped into: under 5 years, 5-14 years (representing children), 15-49 years (representing reproductive age), and 49 years and above.

Insurance status of the patient was defined as having a valid national health insurance card on the day of interview. The types of health care providers that patients sought care from were grouped into three categories: formal health facilities (comprising hospital, clinics, health centres and CHPS), pharmacy shops/licenced chemical shops, and home-based care (left over drugs and traditional care).

The wealth of the household was measured in terms of household assets and possessions using principle component analysis (PCA). We did not collect data on household assets and possessions in our study, but rather used data collected by the Navrongo Health and Demographic Health Surveillance System (NHDSS). The NHDSS database has information on all households in the study district[8]. The study respondent’s perm IDs were linked to the NHDSS data to generate the household wealth index using PCA technique.

A total of 53 households could not be linked with the Navrongo HDSS household assets database and therefore could not be classified and included in the analysis. Thus, household wealth of 340 households were generated. The assets and possessions used for the construction of the wealth index comprised of the type of material used for the wall of the household, roofing material, cooking utensils, toilet facility, source of drinking water and cooking fuel. Household possessions included bicycle, motorbike, car, radio, bed, sewing machine, tape player, television, Digital Versatile Disc (DVD), mobile phone, refrigerator, cattle, sheep, goat, pig and donkey. Households were then assigned to five quintiles: least poor, less poor, poor, very poor, and poorest.

The total cost of fever/malaria per household was calculated by summing the direct and indirect costs incurred by households. The direct cost included both direct medical cost and non-medical costs. The direct medical costs covered all out-of-pocket payments (OOP) for registration card, consultation, diagnosis, medicines and medical supplies on the patient; and direct non-medical costs included all out-of-pocket payments for transportation to and from health facilities (patient and caregiver) and special foods for patients.

Indirect cost (productivity lost) was estimated by multiplying the number of days lost due to malaria by the daily wage for year 2015 (GH¢6/US$1.6). This was calculated for both the patient and the caretaker who were actively engaged in an income-generating activity prior to the illness. In-patient or admission was defined as those who were detained in a health facility for malaria for longer than 12 hours. In-patient costs were therefore the sum of medical costs, laboratory costs, and bed costs during the admission. All costs were collected in Ghana cedis (GH¢) and results presented in both Ghana cedis and US$. The conversion of Ghana cedis to US$ was based on the average exchange rate for 2015 at US$1=GH¢3.8.

### Ethical consideration

Ethical approval for the study was obtained from the Navrongo Health Research Centre Institutional Review Board and the Ghana Health Service Ethics Committee. Written informed consent was obtained from all adult participants and caregivers/parents of children before interviews were conducted. In addition, assent was sought from participants who were between 12-17 years.

## Results

### Background characteristics

Table 1 shows the background characteristics of respondents. Out of the 1,845 households visited, 21.3% (393/1,845) reported to have had an episode of fever or malaria in the past two weeks. The average age of fever patients was 25years and majority of the patients 45% (177/393) were above 49 years old and 26% were under-five years of age. Majority of the fever cases were females 51.7% (203/393) compared to 48.4% (190/393) males. About 83.2% (327/393) of patients had valid national health insurance scheme (NHIS) cards at the time of illness. About 32.3% (127/393) of patients were actively engaged in income generation activity. Out of those who were actively engaged in income generation activity, majority 40.2% (51/127) were farmers, 37.8% (48/127) were artisans, and 22.1% (28/127) were public servants. Most patients 42.8% (168/393) were from poor households (lowest two quintiles).

**Table 1:** Background characteristics of patients who reported fever/Malaria.

| Characteristics | Categories | Number<br>(N=393) | Percent (%) |
| --- | --- | --- | --- |
| Age | Under-five | 102 | 26.0 |
|  | 5 -14 years | 65 | 16.5 |
|  | 15-49 | 49 | 12.47 |
|  | Above 49 | 177 | 45.02 |
| Sex | Male | 190 | 48.35 |
|  | Female | 203 | 51.65 |
| Insurance status | Insured | 327 | 83.21 |
|  | Uninsured | 66 | 16.79 |
| Actively engaged in income generating activity | Yes | 127 | 32.32 |
|  | No | 266 | 67.68 |
| Occupation of those actively engaged in income generating activity | Farmer | 51 | 40.15 |
|  | Public service | 28 | 22.05 |
|  | Artisan | 48 | 37.8 |
| Household wealth | Poorest | 82 | 20.9 |
|  | Very poor | 86 | 21.9 |
|  | Poor/middle | 46 | 11.7 |
|  | Less poor | 52 | 13.2 |
|  | Least poor | 74 | 18.8 |
|  | Missing | 53 | 13.5 |

### Malaria diagnoses and treatment seeking behaviours

Out of the 393 fever patients interviewed, 66.9% (263/393) reported taking antimalarial (ACT). Most of those who used antimalarials 53.6% (141/263) obtained it from formal health facilities (hospitals, Health centres, clinics and CHPS), 41.4 % (109/263) from pharmacy or chemical shops and 5% (13/263) at home. With regards to the type of antimalarial patients used, 36.1% (95/263) reported to have taken Dihydroartemisinin-piperaquine, 35.4% (93/263) took Artesunate–Amodiaquine, 21.7% (57/263) took Artemether-Lumefantrine. Only 49% (195/393) of the study participants had their blood sample taken for illness (microscopic or RDT) and subsequently 73.3% (143/195) were confirmed to have malaria (Table 2).

**Table 2:** Malaria diagnoses and treatment.

| Variables | Categories | Frequency (n=393) | Percent (%) |
| --- | --- | --- | --- |
| Type of treatment | Antimalarial | 263 | 66.9 |
|  | Other treatments | 130 | 33.1 |
| Blood sample taken | Yes | 195 | 49.2 |
|  | No | 198 | 50.8 |
| Confirmed malaria | Yes | 143 | 73.3 |
|  | No | 52 | 26.7 |
| Place where ACT was obtained | Health facility | 141 | 53.6 |
|  | Licensed chemical shop | 109 | 41.4 |
|  | Home | 13 | 5 |
| Type of Antimalarial used | Artesunate–Amodiaquine | 93 | 35.4 |
|  | Artemether-Lumefantrine | 57 | 21.7 |
|  | Dihydroartemisinin-Piperaquine | 95 | 36.1 |
|  | Other Antimalarials (single dose) | 18 | 6.8 |

With regards to prompt treatment, majority of patients did not take antimalarial within 24 hours. For instance, among those who took antimalarial, only 23.6% (62/263) took antimalarial within 24 hours, whiles 44.5% (117/263) took antimalarial between 25 to 48 hours and 31.9% (84/117) took antimalarial after 48 hours of onset of malaria (Table 3).

**Table 3:**
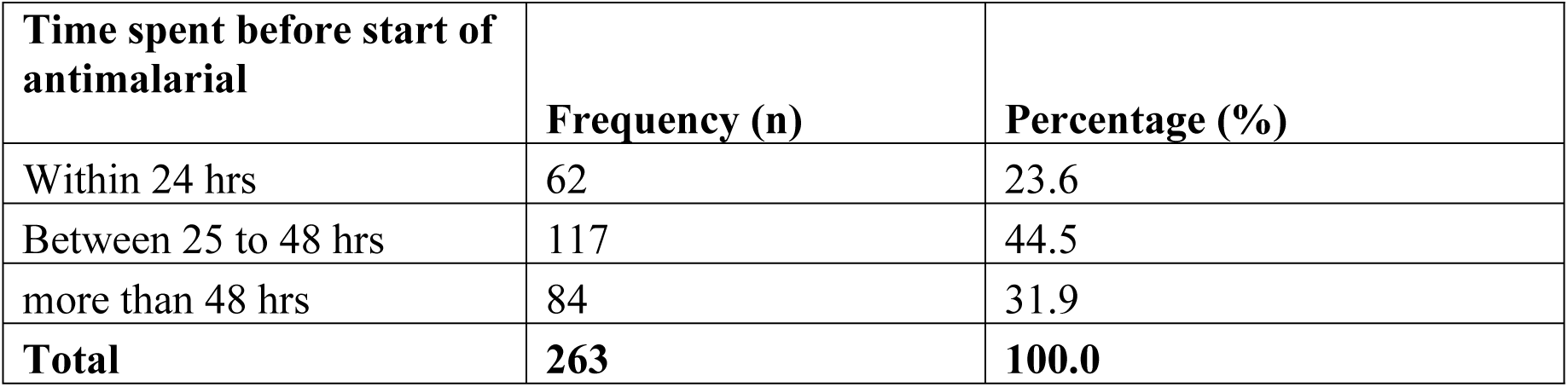
Time spent between onset of fever and start of antimalarial treatment.

### Cost of treating malaria

Table 4 presents direct and indirect costs of treating malaria. Out of the 393 patients, 25.5% (140/393) patients incurred direct medical costs during malaria treatment. The cost was calculated for only those who incurred cost (OOP)in treating the malaria. The average direct medical cost was GH¢11.91 (US$3.13) per treatment. Out of the 140 patients that incurred direct medical costs, 74.3% (104/140) were insured and 25.7% (36/140) were not insured. The average direct medical cost by the uninsured was higher (GH¢14.91/US$3.92) than the insured (GH¢10.87/US$2.86). Out of the total of 393 patients, 82.2% (323/393) spent on direct non-medical costs during the malaria treatment. The direct non-medical coats included transportation and other costs such as special food, water and lodging. About 77.4% (304/393) of respondents incurred transportation cost. The average transportation cost to and from health care providers was estimated at GH¢10.40 (US$2.74). The average direct non-medical costs (transportation and other costs such as special food, water and lodging) was GH¢11.69 (US$3.08). The average direct cost representing medical and non-medical cost was calculated as GH¢14.87/US$3.91 (range: GH¢0.2/US$0.05 - GH¢166/US$43.68). Average indirect cost which relates to productivity lost was estimated at GH¢25.04 (US$6.59). The overall average costs (direct and indirect cost) incurred by households per malaria treatment was GH¢27.82/US$7.32 (range GH¢0.2/US$0.05 - GH¢200/US$52.63).

**Table 4:** Direct and indirect costs of malaria outpatient care (GH¢)

| Cost category | Sub cost Category | Number that paid OOP | Sum total GH¢(US\$) | Mean GH¢(US\$) | Min GH¢(US\$) | Max GH¢(US\$) |
| --- | --- | --- | --- | --- | --- | --- |
| Direct cost | Non-medical cost | 323 | 3775.19(993.47) | 11.69(3.08) | 0.2(0.05) | 108(28.42) |
|  | Medical cost | 140 | 1667.2(438.74) | 11.91(3.13) | 0.5(0.13) | 140(36.84) |
| Total direct cost | Non-medical plus medical cost | 366 | 5442.39(1432.21) | 14.87(3.91) | 0.2(0.05) | 166(43.68) |
| Indirect cost | Indirect cost | 207 | 5184(1364.21) | 25.04(6.59) | 6(0.16) | 114(30) |
| **Overall cost** | **Direct plus indirect cost** | **382** | **10626.39(2796.42)** | **27.82(7.32)** | **0.2(0.05)** | **200(52.63)** |

As presented in Table 5, the average cost incurred in the treatment of malaria by the uninsured was greater (GH¢43.95/US$11.57) than their insured counterparts (GH¢24.75/US$6.51).

**Table 5:**
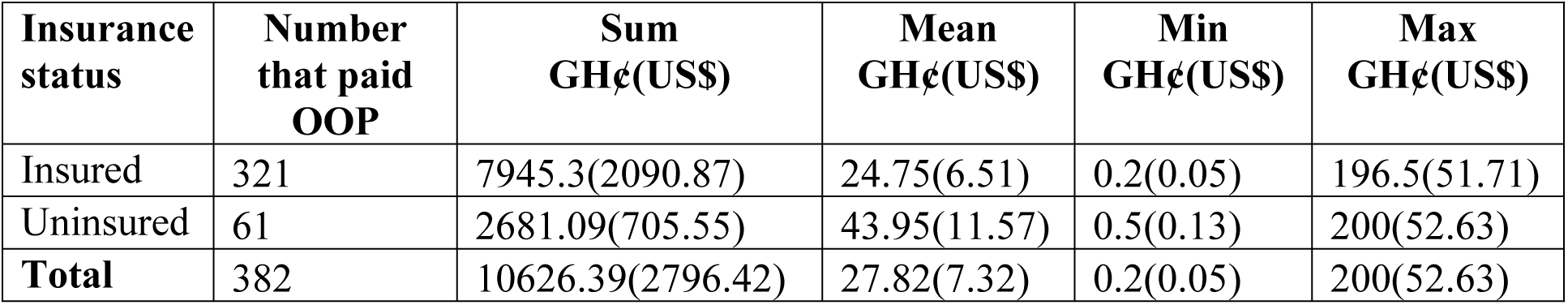
Cost by insurance status.

| Insurance status | Number that paid OOP | Sum GH¢(US\$) | Mean GH¢(US\$) | Min GH¢(US\$) | Max GH¢(US\$) |
| --- | --- | --- | --- | --- | --- |
| Insured | 321 | 7945.3(2090.87) | 24.75(6.51) | 0.2(0.05) | 196.5(51.71) |
| Uninsured | 61 | 2681.09(705.55) | 43.95(11.57) | 0.5(0.13) | 200(52.63) |
| **Total** | **382** | **10626.39(2796.42)** | **27.82(7.32)** | **0.2(0.05)** | **200(52.63)** |

### Hospitalization

Out of the 393 patients interviewed, 13.2% (52/393) were admitted and incurred cost. The average number of days spent by patients that were admitted was 4 days (range 2-7days). The average costs incurred by those who were admitted was GH¢18.94 /US$4.98(range: GH¢0.5/US$0.13 - GH¢150/US$39.47). Surprisingly, the insured paid more hospitalization cost (GH¢20.41/US$5.37) than the uninsured (GH¢14.55/US$3.83).

### Number of days lost due to malaria

The average number of days a malaria episode prevented patients from working was about 5 days for patients and 4 days for the caregivers. Controlling for the patients actively engaged in an income generating activities, the number of days lost still remained at 5 days. The number of school days lost by patients attending school due to malaria was 3 days (range: 1-7 days).

## Discussion

The main aim of the study was to examine the health seeking behaviours as well as cost of treatment of malaria to households in the Kassena-Nankana district of Ghana in the face of multiple first line antimalarial drugs.

This study revealed that a large proportion of malaria patients treated with ACTs (66.9%) and a little more than 50% obtained the ACTs from formal health facilities (hospitals, Health centres, clinics and CHPS). High utilization of formal health facilities for malaria treatment has been reported in a previous studies[11–15]. The high use of formal health facilities in our study could be attributed to NHIS as about 83% of patients were enrolled into the scheme. The positive effects of health insurance enrolment on the utilization of formal health facilities has been widely reported[12,16–19]. Although in our study, there was high utilization of formal health facilities for ACT, buying of ACTs at the pharmacy/chemicals shops was also quite high (41.4%%) and worrisome given that most of the pharmacy/licenced chemical shops do not carry out any diagnostics test before treatment. In other studies conducted in Ghana, however, most people reported visiting pharmacies and chemical shops to treat malaria than the formal health facilities [20–22]. Some of the possible reasons for patient’s preference for chemical shops could be due to waiting time at the health facilities and proximity[20].

Confirmation of malaria infection directs care to those in need. WHO recommends that all persons from endemic areas with suspected malaria should be examined for evidence of infections with malaria parasites by RDT or microscopy[1]. In this regard, the Roll Back Malaria (RBM) partnership had set up a target that at least 80% of malaria patients are diagnosed and treated with effective antimalarial medicines within 24 hours of the onset of illness[23]. Prompt diagnosis and timely treatment can ease illness progression to severe stages and consequently decrease mortality[24]. Given that in our study less than half of patients (49%) had diagnostic tests (Rapid Diagnostic Tests or microscopy) to confirm malaria and only 23.6% took antimalarial within 24 hours of onset of illness is not encouraging and suggests that more efforts are still needed to achieve the RBM target. Jima et al also reported a significant low percentage of respondents who sought prompt care in Ethiopia[25]. People may delay before seeking care because of financial constraint or because they want to be sure they are experiencing symptoms of malaria before they seek care. A possible reason for the low diagnostic test before treatment could be due to limited supply of RDTs at the health facilities[26] where majority of the study participants (53.6%) sought care from.

Some of the reasons for the treatment failure of chloroquine included non-compliance with duration of treatment regimen and misdiagnosis [27,28]. Therefore, diagnosis and rational use of ACTs is crucial in the efforts to reduce malaria burden and to restrain the emergence of ACTs resistance and treatment failure in the near future.

It is interesting to know that despite the fact that Artesunate-Amodiaquine (AA) and Artemether-Lumefantrine (AL) are the most stocked ACTs in the market[29], this study revealed that the most used ACT by patients in the Kassena Nankana District was Dihydroartemisinin-piperaquine (36.1%) followed by Artesunate-Amodiaquine (35.4%) and Artemether-Lumefantrine (21.7%). In fact, in terms of cost of antimalarials, Dihydroartemisinin-piperaquine is the most expensive ACT sold in both the health facilities and the chemical shops in the district. For instance, whereas Artesunate-Amodiaquine(AA) cost about GH¢5/US$1.3 in the pharmacies/chemical shops, P-Alaxin (Dihydroartemisinin-piperaquine) cost about GH¢10/US$2.6, which is twice the price of AA. Contrary to our findings, a previous qualitative study conducted in the same study area reported that respondents preferred artemether-lumefantrine (AL) to other ACTs because AL was perceived to be more effective and has minimal side effects as compared to the other ACTs especially Artesunate-Amodiaquine[30,31]. In our study, a possible reason for the high reported usage of Dihydroartemisinin-Piperaquine could be that patients were interested in trying a new drug and also avoiding the side effects of AA. It also suggests that patients preferred Dihydroartemisinin-Piperaquine because it has the best efficacy and profile in treating uncomplicated malaria as observed in other studies [32]. Our study finding therefore supports the importance of treatment options to take care of those who cannot bear AA and AL.

The overall average cost of treating malaria in the Kassena Nankana District was estimated at US GH¢27.83/US$7.2. Though the cost of treatment is lower than malaria treatment cost estimated in other similar studies conducted in other parts of Ghana [11,33], this amount is still relatively high given that most of the respondents were from poor households. High treatment costs can prevent the poor from seeking care when they have a fever episode as well as impoverish households who seek care when they have an episode of fever [11,34].

The NHIS covers the treatment of about 95% of cases in Ghana including malaria. The benefit package covers both inpatient and outpatient care, emergency and transfer services [35] and the insured are generally not supposed to incur direct medical cost in treating malaria at any formal health facility. In this study, the uninsured incurred more direct cost (GH¢14.91/US$3.9) than the insured (GH¢10.87/US$2.9). Overall, the average cost incurred in the treatment of malaria by the uninsured was greater (GH¢43.95/US$11.57) than their insured counterparts (GH¢24.75/US$6.51). Thus, the uninsured paid US$5 more than the insured, which is consistent with many findings where the uninsured paid more than the insured in treating malaria [12,15,36]. Though the insured were not supposed to incur direct medical costs, a possible reason for the cost incurred by the insured could be due to co-payments for drugs, laboratory test, ward dues and other informal payments. It could also be that antimalarial drugs were out of stock at the health facilities and patient bought drugs outside the health facilities.

The average indirect cost (GH¢25.04/US$6.59) was 1.7 times higher than the direct cost (GH¢14.87/US$3.91). This is consistent with many findings where the indirect costs of malaria treatment were higher than the direct costs of treatment [11,15,33,37,38]. Therefore, efforts to improve access to health care and reduction of financial burden to households in the treatment of malaria should not only be directed to direct medical costs but also indirect costs.

## Limitations

Recall bias is a possible limitation as respondents were asked to recall expenditure over a two weeks period. The reported expenditure could either be overestimated or underestimated especially in cases that respondents did not show receipts on expenditure but reported expenditure verbally. However, because the recall period was short, we do not expect much recall issues to adversely affect the study results.

In the study, we used self-reported fever or malaria to indicate malaria and we acknowledge that as a limitation. For instance, not all fevers are malaria and also it is possible that some reported malaria cases were really not malaria cases. Nevertheless, given that malaria is endemic in the study area and mostly fever is associated with malaria, we assumed high knowledge about malaria signs and symptoms by respondents would not affect the study findings much.

## Conclusions

Though access to antimalarial medicine was quite good, prompt treatment such as treatment within 24 hours of onset of malaria as well as diagnostic testing before treatment of suspected fevers is not encouraging when compared to the 80% target set by the Roll Back Malaria programme. There was a positive effect of national health scheme enrolment on cost of treatment as the insured incurred less cost (US$5 less) in treatment than the uninsured. Health facilities need to be adequately equipped with malaria diagnostic equipment such as Rapid Diagnostic Tests (RDTs) or microscopes and be encouraged to use them so as to direct limited and subsidized antimalarial medicines to patients with malaria parasite infections. Also, more efforts are needed to improve NHIS enrolment in order to reduce cost of treatment and accelerate universal health coverage.

List of abbreviations Competing interests
AA: Artesunate–Amodiaquine
ACTs: Artemisinin-based Combination Therapies
AL: Artemether-Lumefantrine
CHPS: Community Based Health Planning and Services
DP: Dihydroartemisinin-Piperaquine
DVD: Digital Versatile Disc
HDSS: Health and Demographic Surveillance System
INESS: INDEPTH-Network Effectiveness and Safety Studies
IPTp: Intermittent Preventive Treatment for pregnant women
IRS: Indoor Residual Spraying
KND: Kassena-Nankana District
LLINs: long-lasting insecticidal nets
NHIS: National Health Insurance Scheme
NHRC: Navrongo Health Research Centre
OOP: Out-Of-Pocket payment
PCA: Principle Component Analysis
RBM: Roll Back Malaria
RDT: Rapid Diagnostic Test
WHO: World Health Organization

## Competing interests

The authors declare that there are no competing interests.

## Authors’ contribution

MAD PA PDA PAD SC TA and AO: contributed in the conception, design, data collection, analysis, interpretation, and drafting of the manuscript. All authors read and approved the final manuscript.

## Consent for publication

Not applicable

## Availability of data and materials

Relevant data based on which conclusions were made are included in the document. The questionnaire is included as supplementary material.

## Funding

This research was funded by the Bill and Melinda Gate foundation and facilitated by the INDEPTH Network

## Acknowledgement

We wish to express our gratitude to the Navrongo Health Research Centre for the institutional support to undertake the study. We are grateful to the study respondents, data collectors, data managers and the data entry clerks for the various support they provided to this study.

